# Death-associated protein kinase 3 (DAPK3) contributes to intestinal epithelial wound healing and the resolution of experimental colitis in mice

**DOI:** 10.1101/2021.12.03.471118

**Authors:** Huey-Miin Chen, David A. Carlson, Timothy A.J. Haystead, Justin A. MacDonald

## Abstract

Various signaling molecules affecting epithelial restitution and wound healing are dysregulated in ulcerative colitis. Recent evidence demonstrates the necessity of Hippo-YAP/TAZ signaling, interceded by cytoskeletal remodeling, for intestinal regeneration. Death-associated protein kinase 3 (DAPK3) is a regulator of actin cytoskeleton reorganization that controls proliferation and apoptosis. Pharmacological inhibition of DAPK3 in Caco-2 human intestinal epithelial cells (IECs) with the HS38 compound augmented cell proliferation and enhanced wound closure. This phenotype corresponded with the increased colocalization of Yes-associated protein (YAP) with F-actin, which is indicative of YAP activation. The administration of HS38 impeded the resolution of intestinal injury and attenuated epithelial-specific proliferation after acute colitis induced by dextran-sodium-sulphate (DSS) in mice. During recovery from DSS-induced colitis, IEC proliferation was repressed, and mice exhibited increased disease severity when HS38 was applied to inhibit DAPK3. Moreover, HS38 treatment increased YAP nuclear localization in IECs, an indicator of signal activation. In summary, this study established DAPK3 as a key factor in intestinal epithelial regeneration and colitis progression by way of YAP signaling. Nevertheless, the role that DAPK3 play in different cell types will need further investigation to decipher the full consequence of DAPK3 inhibition on epithelial homeostasis.

## 1. INTRODUCTION

Intestinal epithelial cells (IECs) that line the gastrointestinal tract provide a physical and biochemical barrier that separate the host’s internal milieu from foreign antigens and microbial pathogens present in the lumen (Kagnoff, 2014; Pastorelli *et al*, 2013; Peterson & Artis, 2014; Sanchez de Medina *et al*, 2014). Moreover, the epithelium serves as a communication hub for the intestinal mucosa by disseminating luminal events to the sub-epithelial immune compartment and participating in the coordination of mucosal defense and homeostasis (Kagnoff, 2014; Sanchez de Medina *et al*, 2014). Not surprisingly, altered IEC function is frequently detected in ulcerative colitis (UC) patients (Buning *et al*, 2012; Chang *et al*, 2017; Haberman *et al*, 2019; Issenman *et al*, 1993), and has been proposed as a potential etiologic factor for UC (Pastorelli *et al*, 2013). Epithelial integrity may be provisionally weakened by physiological inflammation or minor trauma (Hageman *et al*, 2020; Luissint *et al*, 2016). But, as the balance between pro- and anti-inflammatory stimuli is perpetually disturbed in UC (Strober & Fuss, 2011), epithelial regeneration may be altered so that normal epithelial barrier function is not restored. Inefficacious wound healing can give rise to altered epithelial architecture, impaired barrier function, and intestinal neoplasia which further aggravate the inflammatory state (Koch & Nusrat, 2012; Leoni *et al*, 2015). Therefore, rapid, and complete epithelial wound healing is essential for intestinal health and resolution of UC.

Various signaling molecules integral to epithelial restitution and wound healing are dysregulated in UC; examples include among others: 1) elevated expression of Wnt5a, a noncanonical Wingless-related integration site (Wnt) ligand that facilitates crypt regeneration in wound-associated epithelia (Miyoshi *et al*, 2012) and paradoxically imparts susceptibility to experimental murine colitis (Sato *et al*, 2015); 2) increased production of transforming growth factor-β1 (TGF-β1), an anti-inflammatory cytokine and inducer of tight junction protein and collagen expression (Del Zotto *et al*, 2003; Howe *et al*, 2005; Nigdelioglu *et al*, 2016) that has diminished effect on IECs isolated from IBD patients, partly due to the concurrent increase of TGF-β1-inhibitory molecule Smad7 (Monteleone *et al*, 2004); and 3) constitutive activation of signal transducer and activator of transcription (STAT)-3, a pro-proliferative, anti-apoptotic factor essential for the regeneration of epithelium in response to injury that perpetuates and worsens UC despite its protective attributes (Nguyen *et al*, 2015).

In recent years, research has brought forward the Hippo pathway as a potential therapeutic target for UC. Dysregulation of the Hippo pathway was found in UC patients as well as multiple models of experimental colitis (Deng *et al*, 2018; Nterma *et al*, 2020; Xie *et al*, 2021). Evidence suggests that the major effectors of Hippo pathway, the Yes-associated protein (YAP) and the transcriptional co-activator with PDZ-binding motif (TAZ), link inflammation to epithelial regeneration through their integration of mechanical cues with growth factor signaling (Piccolo *et al*, 2014). Notably, crosstalk between the Hippo/Wnt and the TGF-β1/STAT3 pathways (Gruber *et al*, 2016; Park *et al*, 2015; Qin *et al*, 2018) exposes an inherently complex role for Hippo signaling in intestinal regeneration that requires further investigation in UC (Xie *et al*, 2021).

Death-associated protein kinase-3 (DAPK3) was also recently identified to be a potential key factor in UC progression (Chen & MacDonald, 2021). DAPK3 contributes to cell growth in colon carcinoma cell lines (Togi *et al*, 2011), as well as autophagy induction (Li *et al*, 2021) and wound closure in gastric carcinoma cell lines (Li *et al*, 2015). Furthermore, the reduction of focal adhesion kinase (FAK) activity found with ectopic expression of DAPK3 (Nehru *et al*, 2013), suggests the kinase may also contribute to adhesion regulation during epithelial wound repair, to influence IEC proliferation and survival during UC progression (Owen *et al*, 2011). However, the significance of DAPK3 within these contexts has yet to be directly assessed in experimental models of UC.

While the impairment of several conserved signaling pathways is associated with intestinal epithelial dysfunction in UC (Koch & Nusrat, 2012; Leoni *et al*, 2015; Xie *et al*, 2021), the intricacies of many effector networks involved in the restoration of epithelial homeostasis after injury and/or inflammation are not fully understood. We investigated whether DAPK3 may play a role in epithelial repair and/or UC disease progression given that DAPK3 has been linked to key signaling pathways that coordinate epithelial regeneration after injury. Central to this investigation is the recent availability of ATP-competitive, small molecule inhibitors of DAPK3 that display high specificity and great potency, while bypassing the chemical liabilities that are inherent to other compounds (Al-Ghabkari *et al*, 2016; Carlson *et al*, 2013; Carlson *et al*, 2018).

## 2. METHODS

### 2.1. Materials

The small molecule inhibitors HS38, HS56 and HS94 were synthesized as described previously (Carlson *et al*, 2013; Carlson *et al*, 2018). The Click-iT EdU Cell Proliferation Kit for Imaging, Alexa Fluor™ 488 (#C10337) and 4’,6-diamidino-2-phenylindole (DAPI; #62248) were purchased from Invitrogen-ThermoFisher Scientific. The Phalloidin-iFluor 647 reagent (#176759) was from Abcam (Cambridge, MA). Multiple antibodies were obtained from Cell Signaling Technology (Danvers, MA), including YAP (D8H1X) XP^®^ rabbit mAb (#14074), phosphor[Ser127]-YAP (D9W2I) rabbit mAb (#13008), phospho[Ser397]-YAP (D1E7Y) rabbit mAb (#13619), YAP/TAZ (D24E4) rabbit mAb (#8418), phospho[Thr35]-MOB1 (D2F10) rabbit mAb (#8699), MOB1 (E1N9D) rabbit mAb (#13730), LATS1 (C66B5) rabbit mAb, (#3477), GAPDH (14C10) rabbit mAb (#2118), Ki-67 (D3B5) rabbit mAb (#12202), MST1 rabbit IgG (#3682) and MST2 rabbit IgG (#3952). DAPK3 rabbit IgGs #2928 and #TA313745 were obtained from Cell Signaling and Origene (Rockville, MD), respectively. Cy3-conjugated AffiniPure donkey anti-rabbit IgG (#711-165-152) was from Jackson ImmunoResearch, and dextran sodium sulphate (DSS, 36- 50 kDa; #0216011090) was from MP Biomedicals (Solon, OH).

### 2.2. Cell Culture

Caco-2 human IECs (ATCC #HTB-37) were grown in Dulbecco’s Modified Eagle Medium (DMEM; #11965, Gibco-ThermoFisher Scientific) supplemented with 10% (v/v) fetal bovine serum (FBS; #F1051, Sigma-Aldrich, St. Louis, MO), 100 U/mL penicillin and 100 μg/mL streptomycin (#15140, Gibco), 1 mM sodium pyruvate (#S8636, Sigma-Aldrich), and 1X MEM Non-Essential Amino Acids (#11140, Gibco). Cells were maintained in a 37 °C humidified incubator with 5% CO2 and were routinely passaged every four days at ∼50% confluence using Trypsin-EDTA (0.25%-0.91 mM; #25200, Gibco) solution. All experiments were performed using cells from passage 41-45.

### 2.3. Wound Closure Assays

Caco-2 cells were seeded into Nunclon™ Delta 96-Well MicroWell plates (#167008, ThermoFisher Scientific) at a density of 1.5 x 10^4^ cells/cm^2^ and were maintained for seven days in complete medium. Under these conditions, cells reached confluence in 5-6 days. On day eight, the monolayers underwent overnight serum starvation in preparation for the wound closure assay. On day nine, circular wounds were made using the WoundMaker tool (Essen BioSciences, Ann Arbor, MI). Wounded monolayers were washed twice with serum-free media to remove cellular debris. The wash media was then replaced with treatment media (HS compounds at various concentrations, or DMSO at 1% (v/v), diluted in complete medium, 150 μL/well). Plates were placed into the IncuCyte live-cell imaging system (Essen BioSciences) and maintained in culture at 37 °C, 5% CO2 while whole-well phase images were taken once every hour for 72 hours. ImageJ (Schindelin *et al*, 2012) was used to batch compile overlay masks for images exported from the IncuCyte ZOOM software. Parameters for each step were set based on stacked images from five random wells per experiment, then applied similarly to the entire experiment. Percentage wound closure was fitted to an exponential plateau model [y = ym – (ym – y0) * exp^-k*x^; where y0 is the starting % wound closure (0%), ym is the maximum % wound closure (100%), and k is the rate constant (h^-1^)].

### 2.4. EdU Cell Proliferation Assay

Caco-2 cells were seeded into LabTekII 2-well chambered coverglass (#155380, ThermoFisher Scientific) pre-coated with 0.01% Poly-L-lysine (#P4707, Sigma-Aldrich), at a density of 1.0 x 10^4^ cells/cm^2^. On day eight, the monolayers underwent overnight serum starvation in preparation for wounding. Four scratch wounds per well were created on day nine using a sterile P200 pipette tip attached to an aspirator. The wounded monolayers were washed twice with DMEM then replaced with treatment media (containing 100 μM HS38 or 1% (v/v) DMSO in complete media, 2 mL/well) for 10- or 22-hour incubation in a 37°C humidified incubator with 5% CO2. The Click-iT EdU Cell Proliferation Kit was used as per the manufacturer’s instructions. Post-incubation, wells were washed with 3% (w/v) BSA then PBS, then counterstained with Hoechst 33342 for 30 minutes. After mounting with ProLong™ Glass Antifade Mountant (#P36980, Invitrogen), images were captured using a Nikon A1R laser scanning confocal microscope. One representative image per wound was saved for subsequent analyses. Image analysis was done using the CellProfiler software (McQuin *et al*, 2018). Briefly, Hoechst stain was used for the identification of individual cells, and the EdU signal was used to mark cells undergoing proliferation. Objects defined by the EdU overlay were required to overlap with objects defined by the Hoechst stain to ensure that all EdU-positive objects were true. To determine the percentage of EdU-positive nuclei, roll-up summation was done on count of EdU-positive and count of nuclei for wounds within a single well.

### 2.5. Immunofluorescence Microscopy

Caco-2 monolayers were rinsed twice with PBS prior to a 10-minute incubation with 4% (v/v) PFA. The fixative was removed with two PBS washes. Then, cells were permeabilized with 0.5% (v/v) Tween-20 PBS for 10 minutes. Tween-20 was washed off with PBS (3X, 5 min/wash). Blocking of non-specific binding was completed with 5% (v/v) normal donkey serum in PBS for one hour at room temperature (RT). The blocking solution was replaced by primary antibody for overnight incubation at 4 °C. The YAP (D8H1X) XP^®^ rabbit mAb (14074, Cell Signaling), diluted 1:100 in PBS (containing 1% (w/v) BSA and 0.3% (v/v) Triton X-100) was used to detect endogenous levels of total YAP protein. Cy3-conjugated AffiniPure donkey anti-rabbit IgG was used as the secondary antibody and was applied to samples at 1:200 dilution in conjunction with the Phalloidin-iFluor 647 Reagent used at 1:1,000 dilution in PBS with 1% (w/v) BSA. Prior to room temperature incubation with secondary antibody and phalloidin (one hour), monolayers were washed three times with PBST (PBS containing 0.1% (v/v) Tween-20). Three more PBST washes followed, after which DAPI (10 μg/mL) was added for a 3-minute-long incubation to mark nuclei. Two final PBS washes were carried out before samples were mounted with ProLong™ Glass Antifade Mountant. Images were taken using the Nikon A1R laser scanning confocal microscope on 40X objective (oil) at Nyquist (xy-plane). For image deconvolution, stacks of seven equidistant (1 μm) z-planes were evaluated. The *Colocalization GUI* toolbox for MATLAB (MathWorks, Natick, MA) was used for image and statistical analysis of YAP-F-actin colocalization (Villalta *et al*, 2011). The *Colocalization GUI* algorithm generates a colocalization mask for the calculation of Manders coefficients *m*1 and *m*2 (Manders *et al*, 1993). The mask represents pixels that display true (i.e., non-random) colocalization in a pair of images. The *m*1 and *m*2 parameters represent the percentage of image 1 (YAP) or image 2 (F-actin) intensity that is colocalized with matched pixels on image 2 (F-actin) or image 1 (YAP), respectively. MAPs are the visual representation of the contribution of each pixel to the colocalization coefficients *m*1 or *m*2. Two *z*-slices per stack (of seven equidistant z-planes) were selected for colocalization analysis based on the broad presence of distinct cortical actin belt across the full image.

### 2.6. Mouse model of dextran sodium sulphate (DSS)-induced colitis

All animal protocols were approved by the Animal Care and Use Committee at the University of Calgary and conform to the guidelines set by the Canadian Council of Animal Care. Experimental UC was induced in nine to 11-weeks old male C57BL/6 mice by the addition of 2.5% (w/v) DSS in drinking water for seven days. For recovery, mice were provided regular drinking water for two days post-DSS- acclimatization (DSS+2); at which time, animals were euthanized by cervical dislocation after anesthesia by isoflurane inhalation. Body weight was tracked daily then normalized to day one of DSS administration. A disease activity index (DAI) score was estimated based on weight, stool consistency, and blood in feces (**Table 1**). Macroscopic scoring of dissected colon tissue was performed according to the grading system outlined in **Table 2**.

**Table 1.** Criteria for the scoring of disease activity index (DAI) for mice subjected to the DSS- associated colitis model.

| Parameter | Score |  |  |  |
| --- | --- | --- | --- | --- |
|  | 0 | 1 | 2 | 3 |
| **Weight Change** | <= 5% | > 5% | > 10% | > 15% |
| **Stool Consistency** | Normal | Loose Stool | Diarrhea |  |
| **Blood in Stool** | Normal | Slight | Gross |  |

**Table 2.** Criteria for the macroscopic scoring of mouse colon tissue dissected from the DSS- associated colitis model.

| Parameter | Score |  |  |  |
| --- | --- | --- | --- | --- |
|  | 0 | 1 | 2 | 3 |
| **Colon Shortening** | <5% | 6-15% | 16-30% | >30% |
| **Colonic Edema** | Normal | >=1 cm | >=3 cm | >=5 cm |
| **Adhesion** | None | Minor | Major |  |
| **Inflammation** | Normal | Localized Hyperemia | Ulceration w/o Hyperemia | Ulceration with Inflammation |

### 2.7. Histological Examination of Epithelial Damage and Inflammation

Paraffin-embedded tissues (distal colon), sectioned at 5-μm thickness, were deparaffinized, rehydrated with gradient ethanol, then stained with hematoxylin and eosin (H&E) for blinded microscopic assessment of colitis severity. The grading system considers the degree of epithelial damage and infiltration of inflammatory cells (Hirota *et al*, 2011), as well as the percentage of involvement (**Table 3**).

**Table 3.** Criteria for the histological scoring of epithelial damage and inflammation in distal colons isolated from the DSS-associated colitis model.

| Parameter | Score |  |  |  |  |
| --- | --- | --- | --- | --- | --- |
|  | 0 | 1 | 2 | 3 | 4 |
| **Epithelial Damage** | Normal | Villus Vacuolation/<br>Bleeding | Loss of Epithelium | Complete Loss of Crypt Architecture |  |
| **% Architectural Change** | 0% | 1-30% | 31-60% | >60% |  |
| **Inflammation** | Normal | Increased Inflammatory Cells in the Lamina Propria | Increased Inflammatory Cells in the Submucosa | Dense Inflammatory Cell Mass | Transmural Inflammation |
| **% Inflammation** | 0% | 1-25% | 26-50% | 51-75% | >75% |

### 2.8. Immunohistochemistry

Deparaffinized tissues were rehydrated with gradient ethanol, then subjected to heat-induced antigen retrieval in 10 mM sodium citrate (pH 6.0). Sections were subsequently washed with PBST, then blocked with 10% normal donkey serum in PBS. Overnight incubation was carried out at 4 °C with anti-Ki-67, anti-DAPK3, or anti-YAP, diluted 1:400 in PBS. The Cy3-conjugated AffiniPure donkey anti-rabbit IgG was the secondary antibody and was applied to samples at 1:200 dilution. DAPI was used at 10 μg/mL to mark nuclei. For sections probed with anti-Ki-67, images were taken with the Olympus FV10i confocal microscope on the 60X objective (oil). Ki-67^+^ positive nuclei was manually counted, then expressed as a percentage of total nuclei per crypt. Between 19-30 well-oriented crypts from 2-3 sections were evaluated per animal. Sample size was 3-8 animals per group. For sections probed with anti-DAPK3, images were taken with the Olympus FV10i confocal microscope using a 60X objective (oil). For sections probed with anti-YAP, images were taken with Nikon A1R laser scanning confocal microscope on the 40X objective (oil). For image deconvolution, stacks of 10 equidistant (0.5 μm) z-planes were evaluated.

### 2.9. Immunoblotting

Midsections from murine colon were homogenized in lysis buffer (20 mM Tris-HCl (pH 8.0), 100 mM NaCl, 0.1% (w/v) SDS, 0.9 mM EDTA, 0.5% (v/v) Triton X-100, 7 mM Na3VO4, 50 mM NaF, and cOmplete™ Protease Inhibitor Cocktail). Homogenates were sonicated for 10 minutes at 4 °C, then centrifuged at 14,000 rpm for 10 minutes. Supernatants were collected, and proteins were resolved by SDS-PAGE (12%) gel electrophoresis then transferred onto 0.2 μm nitrocellulose membranes. Blocking was done with 5% (w/v) non-fat milk in TBST, then membranes were incubated overnight in primary antibodies at 4 °C. Immunoreactive proteins were visualized using the ChemiDoc Imaging System (Bio-Rad Laboratories, Hercules, CA).

### 2.10. Statistical Analyses

Data are presented as mean ± SEM, and the statistical analyses were performed with GraphPad Prism 8.3.0 (GraphPad Software, La Jolla, CA). Unless otherwise noted, the Student’s unpaired t-test was used to compare between 2 groups; one-way ANOVA with Dunnett’s *post hoc* test or two-way ANOVA followed by Tukey’s multiple comparison test was used to assess more than 2 groups. P values <0.05 were considered significant.

## 3. RESULTS

### 3.1 monolayer wound closure is enhanced with DAPK3 inhibitor treatment

As shown in **Figure 1A**, treatment with HS38 significantly enhanced Caco-2 wound closure. At 36 h post-wounding, administration of HS38 (100 μM) increased the wound closure by 46% when compared to the DMSO vehicle control (**Figure 1B**). Because of enhanced wound closure, many wounds treated with HS38 became segmented after the 36 h time point and created multiple (smaller) wound areas in one single well (data not shown). The presence of segmented wounds obstructed the ability to complete effective data processing and analyses. Consequently, the time frame for monitoring the wound closure was reduced to 36 h. **Figure 1C** shows the percentage wound closure over time, overlayed with curves fitted on to an exponential plateau model to evaluate the rate of closure (Arciero & Swigon, 2013; Jonkman *et al*, 2014). The wound closure rate constants support the notion that pharmacological inhibition of DAPK3 with HS38 increases the rate of wound closure in a concentration-dependent manner (**Figure 1D**). Examination of the fitted curves (**Figure 1C**) suggests that at higher [HS38], the rate of wound closure might not be restrained by the parameters set forth for the modeling exercise (i.e., plateau ≠ 100%). To investigate further, the first derivatives of the percentage wound closure curves were calculated. As shown in **Figure 1E**, the derivative for the rate of wound closure decreased over time for the control conditions while the HS38 treatment did not attenuate the rate despite having reached a greater percentage of closure at 36 h. Presumably, DAPK3 inhibition via HS38 interfered with the sensing of chemical and/or mechanical environmental cues that would otherwise inform the collective behaviour of the reparative monolayer.

**Figure 1.**
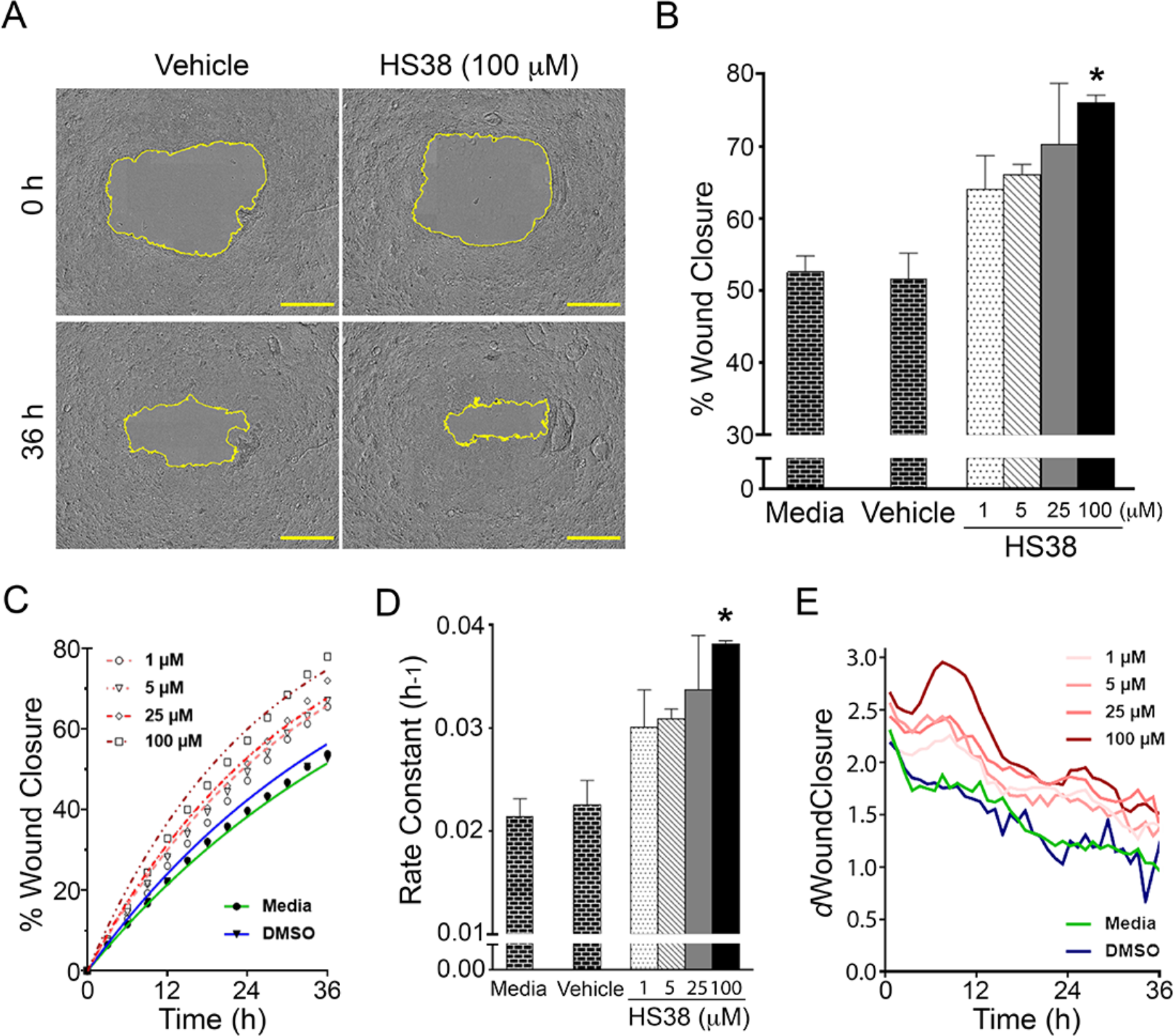
DAPK3 inhibition with HS38 *in vitro* increased Caco-2 wound closure in a concentration-dependent manner. Circular wounds were made on Caco-2 monolayers, and the size of wound areas was measured once per hour as the monolayers healed. Representative images **(A)** are shown for vehicle (DMSO) and HS38 (100 μM) treatments at the 36 h time point. The images are representative of three independent experiments. Scale bars = 800 μm. The percent wound closure at 36 h relative to the initial wound size **(B)**, the percentage wound closure over time with data plotted at 3 h intervals **(C)**, the rate constants (*k*, h^-1^) derived by fitting the percent wound closure curve to an exponential plateau model **(D)**, and the first derivative (*d*WC) of the percent wound closure curve **(E)** are provided for increasing [HS38] (1, 5, 25, or 100 μM). One-way ANOVA with Dunnett’s *post hoc* test comparing the five treatment groups (vehicle and HS38 at various concentrations) indicate significant difference among means, * p<0.05. Error bars represent ± SEM.

In addition to DAPK3, HS38 exhibits high potency towards DAPK1 and the structurally related proviral integrations of moloney virus 3 (PIM3) kinase (**Table 4**; Carlson *et al*, 2013). To differentiate the DAPK3-mediated phenotype from the PIM3-mediated phenotype, parallel experiments were conducted by treating wounded Caco-2 monolayers with HS38, HS56 (more inhibitory activity towards PIM3), or HS94 (less inhibitory activity towards PIM3) (Carlson *et al*, 2018). As illustrated in **Figures 2A** and **2B**, HS94 yielded a similar phenotype as HS38, whereas HS56 treatment produced an opposite phenotype. That is, monolayers treated with HS56 yielded a reduction in percentage wound closure (21% decrease vs. DMSO vehicle control), and monolayers treated with HS94 experienced enhanced wound closure (19% increase vs. DMSO vehicle control) at 36 h. The wound closure of monolayers treated with HS56 plateaued at around 40%. In several independent replicates, wound closure of Caco-2 monolayers treated with HS56 halted at about 36 h post-wounding and failed to achieve 100% wound closure (data not shown). The rate constants derived from modeling exhibited the same trend as what was reflected by the 36 h end-point measurement of percentage wound closure. That is, HS94 yielded a greater rate constant whereas HS56 produced a lower rate constant (**Figure 2C)**. Interestingly, in the early phase of wound closure, treatment with HS56 increased the rate of wound closure (**Figure 2D)**; however, this was not sustained, and a rapid deceleration in wound repair was observed as time advanced.

**Figure 2.**
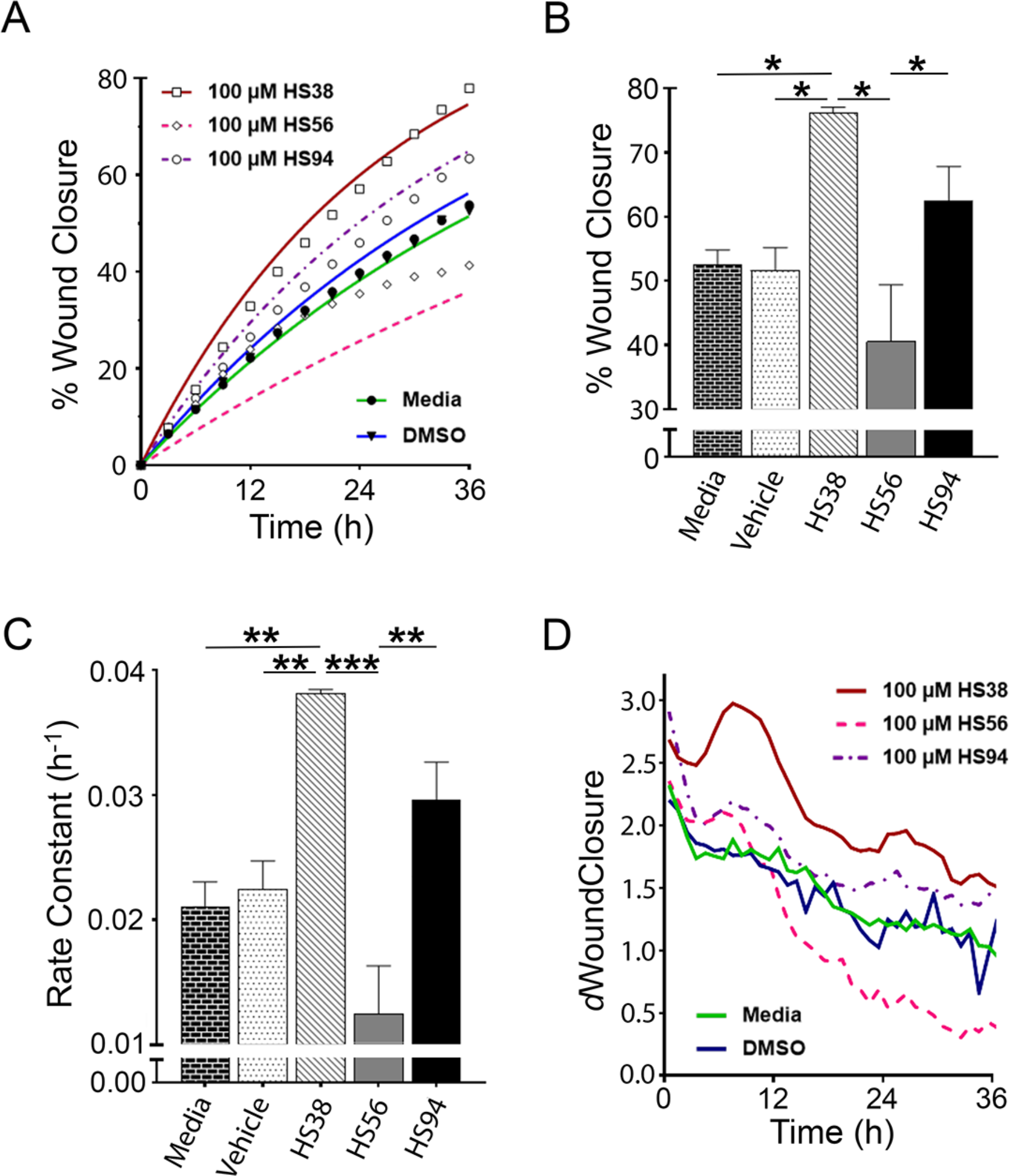
*In vitro* wound closure effects obtained with HS38 can be ascribed to DAPK3 inhibition. Circular wounds were made on Caco-2 monolayers, and the percent wound closure over time was plotted at 3 h intervals **(A)**. The percent wound closure after 36 h was compared for monolayers treated with HS38, HS56 or HS94 (all at 100 μM). Error bars represent ± SEM. **(B)**, the rate constants (*k*, h^-1^) derived by fitting the % wound closure curve to an exponential plateau model **(C)**, and the first derivative (*d*WC) of the percent wound closure curve **(D)** are provided. Error bars represent 95% CI. Figures are representative of three independent experiments.

**Table 4.** Inhibition constants for the HS compounds.

|  | K<sub>i</sub> (μM) |  |  |
| --- | --- | --- | --- |
|  | HS38 | HS56 | HS94 |
| DAPK3 | 0.260 | 0.315 | 0.126 |
| PIM3 | 0.208 | 0.072 | 2.50 |

Data extracted from Carlson *et al.*, 2018.

### 3.2 cell proliferation is augmented with application of the DAPK3 inhibitor HS38

The closure of epithelial wounds involves epithelial cell proliferation and migration. To examine the effects of HS38 administration and DAPK3 inhibition on cellular proliferation in wounded monolayers, EdU-coupled fluorescence was used to evaluate cell proliferation around wound edges at 12- and 24-hours after wounding. In this case, HS38 treatment did not alter the percentage of EdU-positive cells at 12 hours post-wounding (**Figure 3A**). However, at 24 hours post-wounding, cell proliferation in HS38-treated monolayers was significantly greater than in the control monolayer (**Figure 3B**; HS38: 38% EdU-positive, DMSO: 18% EdU-positive; p<0.01). This finding suggests that the enhanced rate of wound closure seen with HS38-treated monolayers (**Figure 1**) was owed, in part, to an augmentation of cell proliferation by DAPK3 inhibition.

**Figure 3.**
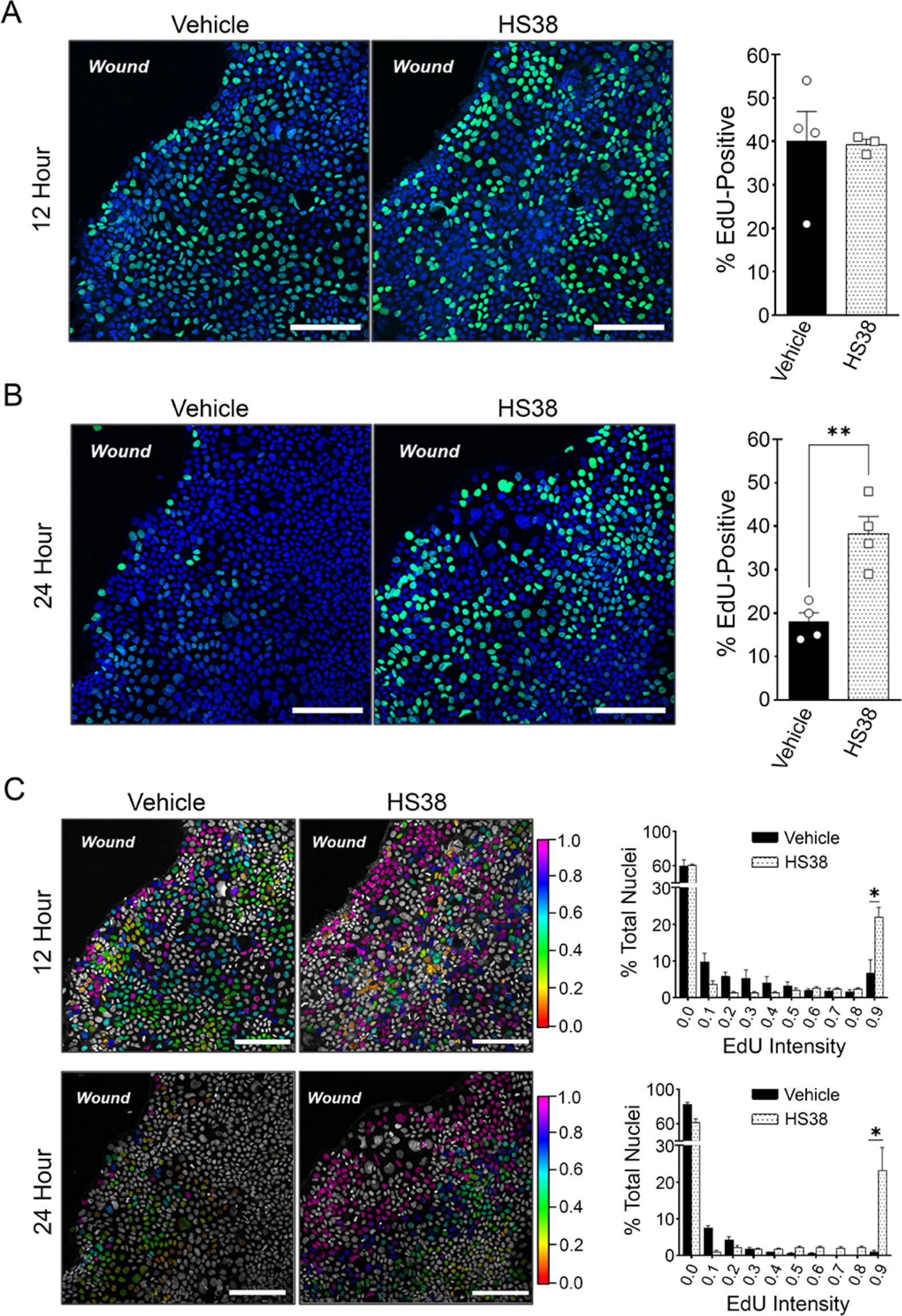
DAPK3 inhibition alters Caco-2 proliferation and cell cycle progression during *in vitro* wound closure. Circular wounds were made on Caco-2 monolayers. EdU-labeled proliferation was assessed after treatment with HS38 at 100 μM for 12 hours **(A)** or 24 hours **(B)**. In **(C)**, the distribution of EdU-coupled fluorescence intensity was determined after 12 hours or 24 hours of vehicle (DMSO, 1%) or HS38 (100 μM) treatment. Statistical significance was determined via unpaired Student’s t-test; * p<0.05, ** p<0.01. Error bars represent ± SEM for n=3-4 independent experiments. Scale bars = 300 μm.

Although the percentage of EdU-positive cells did not change in response to HS38 treatment at the 12 h time point, an interesting observation was made when EdU intensity levels were transformed from a nominal to an interval scale (**Figure 3C**). When cells in the EdU-positive population were partitioned based on relative intensity, it was apparent that most EdU-positive cells in the HS38-treated monolayers exhibited maximal-intensity (>0.9), whereas the opposite was true for cells in the control monolayer (**Figure 3C**; EdU-positive cells showing maximal-intensity: HS38, 56%, Control, 16%; p=0.024). Similar bias was detected for monolayers treated for 24 hours with HS38; among the EdU-positive cells exposed to HS38, 57% were found to have reached maximum intensity, whereas only 5% of the EdU-positive cells (p=0.012) in control monolayers achieved maximum intensity (**Figure 3C)**.

### 3.3 Treatment of Caco-2 monolayers with HS38 increased colocalization of YAP with F-actin

Previous reports have associated DAPK3 expression and/or activity with perturbed cell cycle kinetics (Hu *et al*, 2020; Ono *et al*, 2020; Wu *et al*, 2015) and actomyosin cytoskeletal remodelling (Boosen *et al*, 2009; Komatsu & Ikebe, 2004; Nehru *et al*, 2013). Moreover, mechanical signaling via alterations to the actin cytoskeleton is a critical controller of Yes-associated protein (YAP) (Aragona *et al*, 2013; Wada *et al*, 2011). Studies have shown the colocalization of YAP with F-actin under YAP-activating conditions (Lee *et al*, 2019) as well as reported an association of YAP expression and/or activity with S-phase shortening and promotion of G1/S transition (Cabochette *et al*, 2015; Cai *et al*, 2010). These findings, along with a potential regulatory relationship identified for *DAPK3* and *YAP* in colitis-associated dysplasia (Chen & MacDonald, 2021), provide impetus to complete additional investigations of DAPK3 inhibition on the colocalization of YAP with F-actin.

The colocalization of YAP and F-actin immunofluorescence (**Figure 4A**) was evaluated at the wound edges of Caco-2 monolayers at 24 hours post-wounding by computation with a colocalization mask algorithm (Villalta *et al*., 2011). The *m*1 MAP, providing visual representation of the contribution of each pixel to the colocalization coefficients *m*1, is displayed in **Figure 4B**. The *m*1 coefficient values, which represent the percentage of YAP-positive pixels that are colocalized with matched pixels on the F-actin channel, are summarized in **Figure 4C.** The *m*1 mean value for HS38-treated monolayers was significantly greater than for the control monolayers (+0.253 ± 0.079, p=0.0064). Since *m*1 values represent the percentage of YAP-positive pixels that also display non-random fluorescence in the phalloidin channel, this result suggests that DAPK3 inhibition increased colocalization of YAP with F-actin. Even though the difference in *m*1 value between treatment types was significant, several control monolayers exhibited similar *m*1 values as HS38 treated monolayers (**Figure 4C**). Potentially, the *m*1 quantification result for the control monolayers included data points that were driven by high, but focused, YAP staining at defined subcellular compartment/components. Qualitative analysis of *m*1 MAPs generated for HS38-treated and control monolayers supported this hypothesis. As shown in **Figure 4B**, monolayers treated with HS38 displayed a qualitative difference in dispersion of *m*1 signal intensity throughout the entire cell cytoplasm, whereas the *m*1 signal intensity are localized to the cortical actin ring on the cell periphery in the control monolayer. The visual phenotype displayed in **Figure 4B** is representative of the general trend observed for the two treatment types. One interpretation of these data is that the inhibition of DAPK3 initiated YAP activation via dispersion of YAP from the cell membrane. Continued association of YAP with F-actin in the cytoplasm might aid the transport of YAP from cell periphery to the nuclear membrane. It should be noted, however, that YAP immunofluorescence exhibited few instances of nuclear localization in monolayers treated with either HS38 or vehicle (**Figure 4A**).

**Figure 4.**
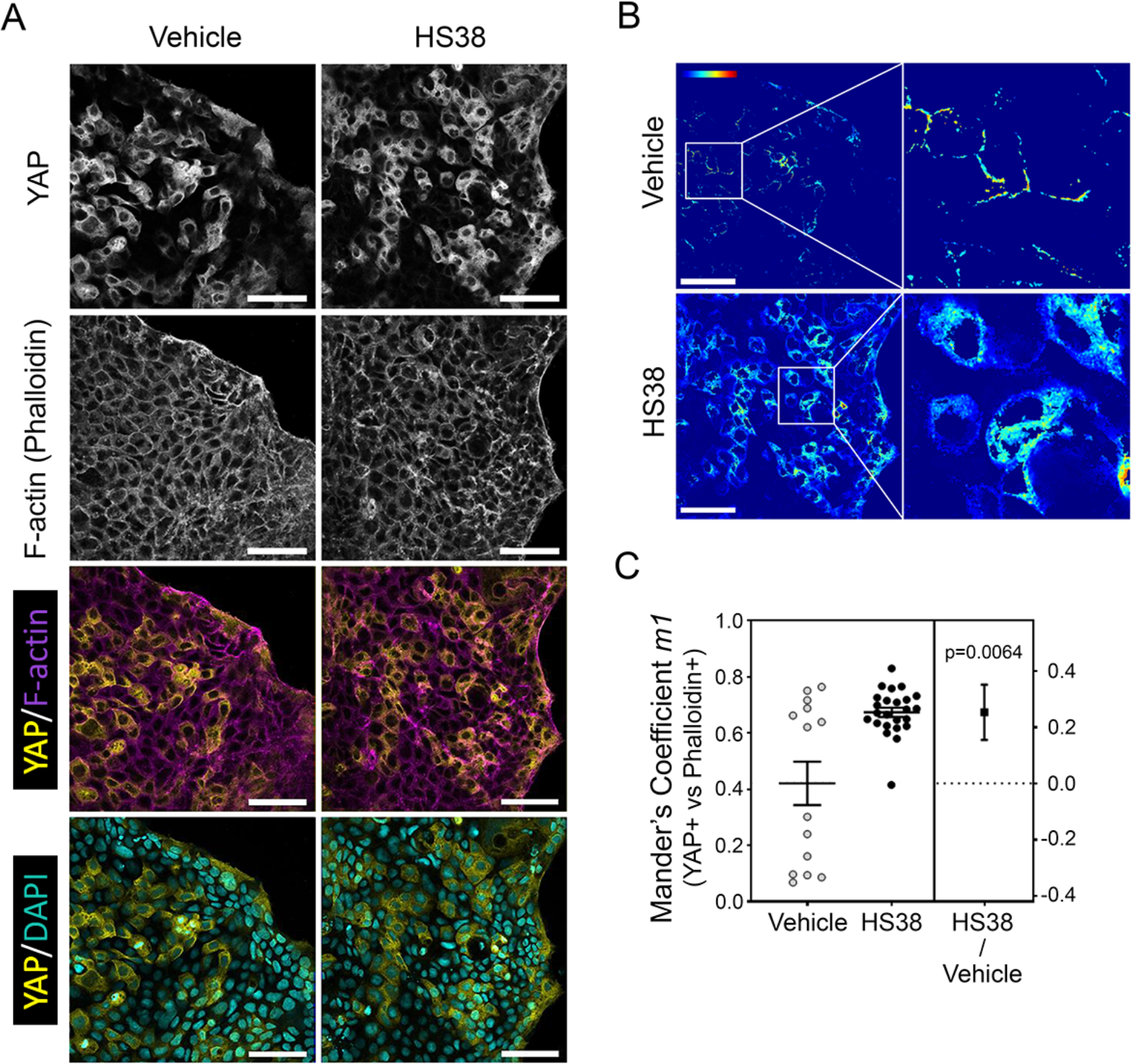
*In vitro* DAPK3 inhibition increased YAP colocalization with F-actin. Immunofluorescence images **(A)** of Caco-2 monolayers were taken with a laser scanning confocal microscope at Nyquist (xy-plane). For image deconvolution, stacks of seven equidistant (1 μm) z-planes were evaluated. In **(B)**, *m*1 MAPs of representative images from the HS38-treated and control groups. Jet color bar: contribution of each pixel to the m*1* coefficient. Scale bars = 300 μm. In **(C)**, comparison of *m*1 coefficient values between HS38-treated (100 μM) and vehicle control (DMSO, 1%) monolayers showed significant increase in colocalization of YAP and phalloidin immunofluorescence. Datapoints represent z-slices analyzed for colocalization; error bars represent ± SEM, n=3-4 independent experiments.

### 3.4 In vivo HS38 administration increased susceptibility to DSS-induced colitis

Mice were administered HS38 by *s.c.* injection and then provided DSS in drinking water (**Figure 5A**). Mice administered DSS and HS38 showed greater weight loss than their vehicle-control littermates (p<0.0001; **Figure 5B**). Moreover, two of the DSS mice that were co-treated with HS38 had to be removed from the study due to severe weight loss that exceeded the humane endpoint criteria (i.e., weight loss of >15%). While there was no general variation in DAI (disease activity index) over the course of DSS administration, *post hoc* multiple comparison showed significant HS38-dependent change in DAI at DSS+2 (i.e., seven days of DSS treatment plus two days of recovery in the presence or absence of HS38 treatment; p=0.039, **Figure 5C**). Colon lengths (**Figure 5D**) and macroscopic scores (**Figure 5E**) exhibited subtle, but non-significant, differences between HS38- and vehicle-treated mice at DSS+2, and histological analyses revealed a small but significant increase in colitis severity for the HS38-treated mice, as per measures of epithelial damage and inflammation (**Figures 5F**, **5G**). Taken together, these data suggest that DAPK3 plays a protective role in DSS-induced colitis.

**Figure 5.**
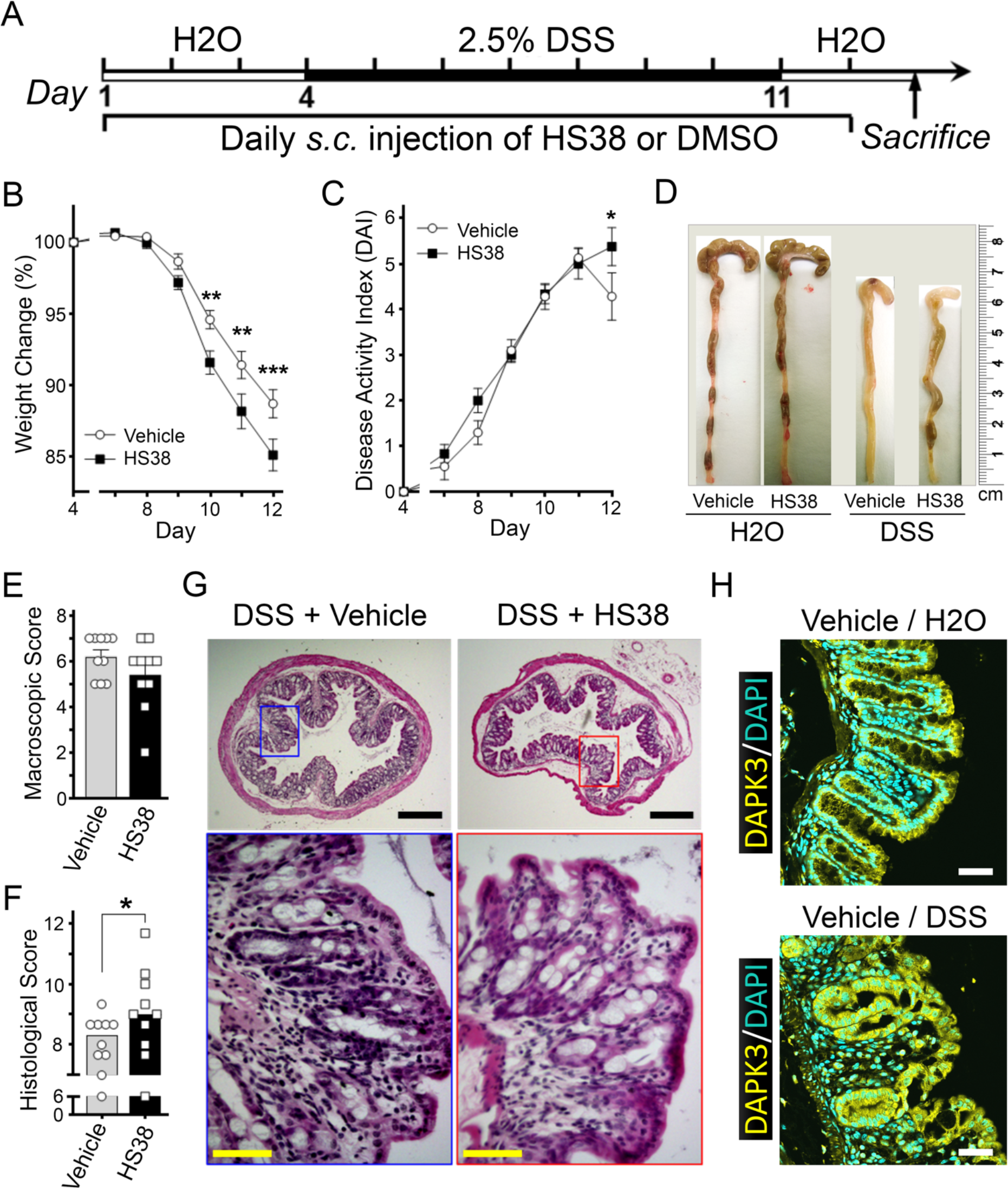
DAPK inhibition with HS38 *in vivo* increased susceptibility to DSS-induced colitis. Mice were subjected to experimental colitis **(A)** by providing 2.5% DSS in drinking water *ad libidum* for 7 days while receiving daily injections of HS38 (500 μg/kg) or vehicle (DMSO). Animals were then sacrificed after two days recovery on normal water (DSS+2). The percentage weight change **(B)** and disease activity index, DAI **(C)** were monitored over the 8 days when mice were co-treated with DSS and HS38 (n=12) or vehicle control (n=10). Statistical significance was determined via multiple comparison analysis after 2 way-ANOVA with the Šídák correction method; ** p<0.01, *** p<0.001. Colons were dissected **(D)** and macroscopic scores **(E)** were determined for HS38- and vehicle-treated mice at DSS+2. Histological scoring **(F)** was completed using H&E-stained transverse sections **(G)** of colons from vehicle- and HS38-treated mice; scale bars (black) = 300 μm, scale bars (yellow) = 50 μm. Statistical significance was determined via unpaired Student’s t- test; * p<0.05. Error bars represent ± SEM. DAPK3 immunoreactivity was identified in the intestinal epithelium of mice **(H)**. Cross-sections of the distal colon were fixed and probed with anti-DAPK3 and counterstained with DAPI. Scale bars, 50 μm.

### 3.5. Increased DAPK3 in IECs is identified with recovery from DSS-induced injury (DSS+7)

Immunohistochemistry was used to probe DAPK3 protein localization within the distal colon of mice **(Figure 5H)**. DAPK3 immunoreactivity was observed for epithelial cells in tissue sections dissected from both healthy (H2O) and colitis (DSS) animals. The colonic epithelium of healthy animals was characterized by punctate DAPK3 immunoreactivity most prominently observed in the cytoplasm of IECs at the villus tips and the crypt base columnar cells. Prominent immunoreactivity was observed for the epithelium of colitis animals with cytoplasmic DAPK3 staining. As expected, villous atrophy and other morphological alterations were apparent in colons of DSS-colitis animals, and demonstrable DAPK3 immunoreactivity was detected in the crypts of the damaged epithelium. Colonic crypts were isolated from distal colon tissue (Magness et al, 2013) to further profile DAPK3 in IECs. The epithelial expression of *Dapk3* was not impacted by DSS-induced colitis or HS38 treatments (**Figure 6A**); however, DAPK3 protein abundance was significantly elevated in IECs isolated from animals in the recovery phase following removal of DSS (**Figure 6B**). Flow cytometry and gating with EpCaM (epithelial), CD31 (endothelial) and CD45 (hematopoietic) confirmed that > 75% of cells in the cellular analytes were epithelial in nature (**Figure 6C**). Taken together, these data support the concept that DAPK3 functions as a responding element within IECs during the resolution of experimental colitis.

**Figure 6.**
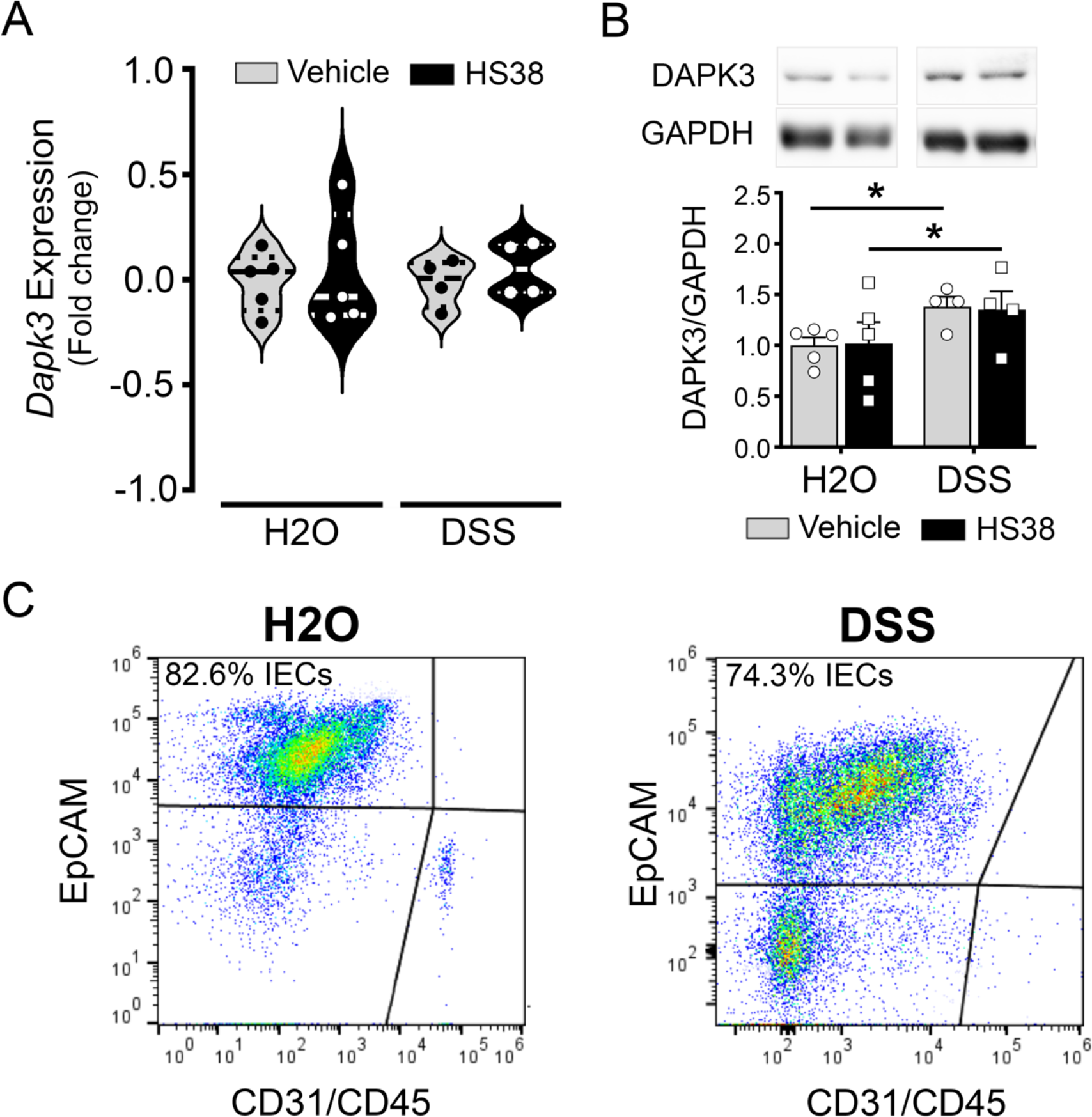
Increased DAPK3 abundance is identified in colonic IECs following recovery from DSS- induced colitis. Mice were subjected to experimental colitis by providing 2.5% DSS in drinking water while receiving HS38 or vehicle (DMSO). Animals were sacrificed after seven days recovery on normal water (DSS+7). Colonic crypts were isolated, then cell lysates were prepared and subjected to qPCR for quantitation of *DAPK3* expression **(A)** or immunoblot analysis for DAPK3 protein abundance **(B)**. Flow cytometry was used to assess the proportion of IECs recovered following isolation of colonic crypts **(C)**. Gating on epithelial cell adhesion molecule (EpCAM)^+^ and CD45^+^/CD31^+^ were used to distinguish IECs from hematopoietic and endothelial cell populations.

### 3.6. In vivo HS38 administration attenuated IEC proliferation post DSS-induced injury

To determine the proliferative fraction of IECs in mice, paraffin-embedded distal-colon tissues were probed with antibodies targeting Ki-67 (**Figure 7A**). Mice given regular drinking water showed no significant differences in IEC proliferation regardless of treatment with HS38 or vehicle (**Figure 7B**). In these animals, Ki-67^+^ cells were displayed at the crypt base and encompassed ∼25% of total crypt IECs (**Figure 7B**). In response to DSS-induced injury, the proportion of Ki-67^+^ cells increased to ∼57% of total IECs per crypt. In mice recovering from DSS-induced injury, HS38 treatment decreased the epithelial distribution of proliferating cells to levels like those of uninjured animals (∼32%, **Figure 7B**). These data indicate that the administration of HS38 impeded the resolution of intestinal injury and attenuated epithelial-specific proliferation after acute colitis induced by DSS.

**Figure 7.**
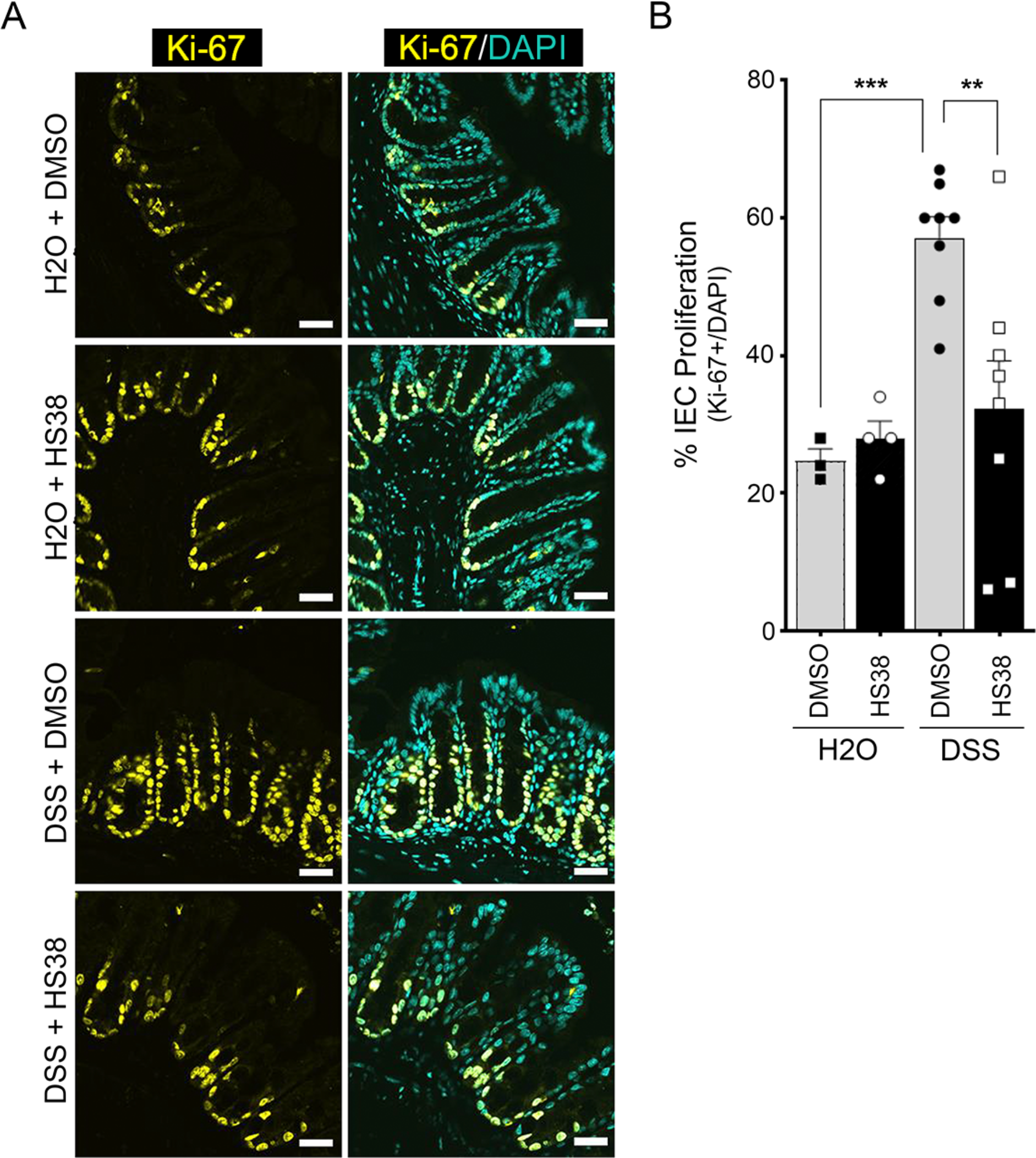
DAPK inhibition attenuated colonic IEC proliferation following DSS-induced injury. **(A)** Representative Ki-67 staining of the colon of HS38-treated and vehicle control mice is provided. Scale bars = 50 μm. **(B)** IEC proliferation was quantified in the colon and is provided as the percentage of Ki-67^+^ cells related to the total DAPI^+^ cells per crypt. Statistical significance was determined by 2-way ANOVA with Tukey’s post hoc multiple comparisons test ** p<0.01, *** p<0.001; n=3-8 animals. Error bars represent ± SEM.

### 3.7. HS38 administration stimulated YAP nuclear accumulation in vivo

The effects of HS38 treatment on the core Hippo signaling pathway and the cellular localization of YAP was investigated since a change in IEC proliferation, brought about by DAPK3 inhibition and cytoskeletal destabilization, may involve the Hippo pathway (Aragona *et al*, 2013). Intriguingly, in >50% of uninjured animals, YAP protein abundance appeared elevated in response to DAPK3 inhibition (**Figure 8A**); however, this difference did not reach statistical significance. DAPK3 inhibition was accompanied with decreased pS127-YAP levels (**Figure 8B**), significantly so in DSS-treated animals but not in control animals. However, there was no significant impact of DAPK3 inhibition on the levels of p397-YAP (**Figure 8C**). None of the core Hippo regulators showed significant changes in protein abundance upon DAPK3 inhibition regardless of injury status, including MST1, MST2, LATS1 as well as MOB1 (**Figure 9A-D**). A decrease in pT35-MOB1 was observed with DSS-induced injury; however, no change in this MOB1 phosphorylation was apparent with HS38 administration (**Figure 9E**).

**Figure 8.**
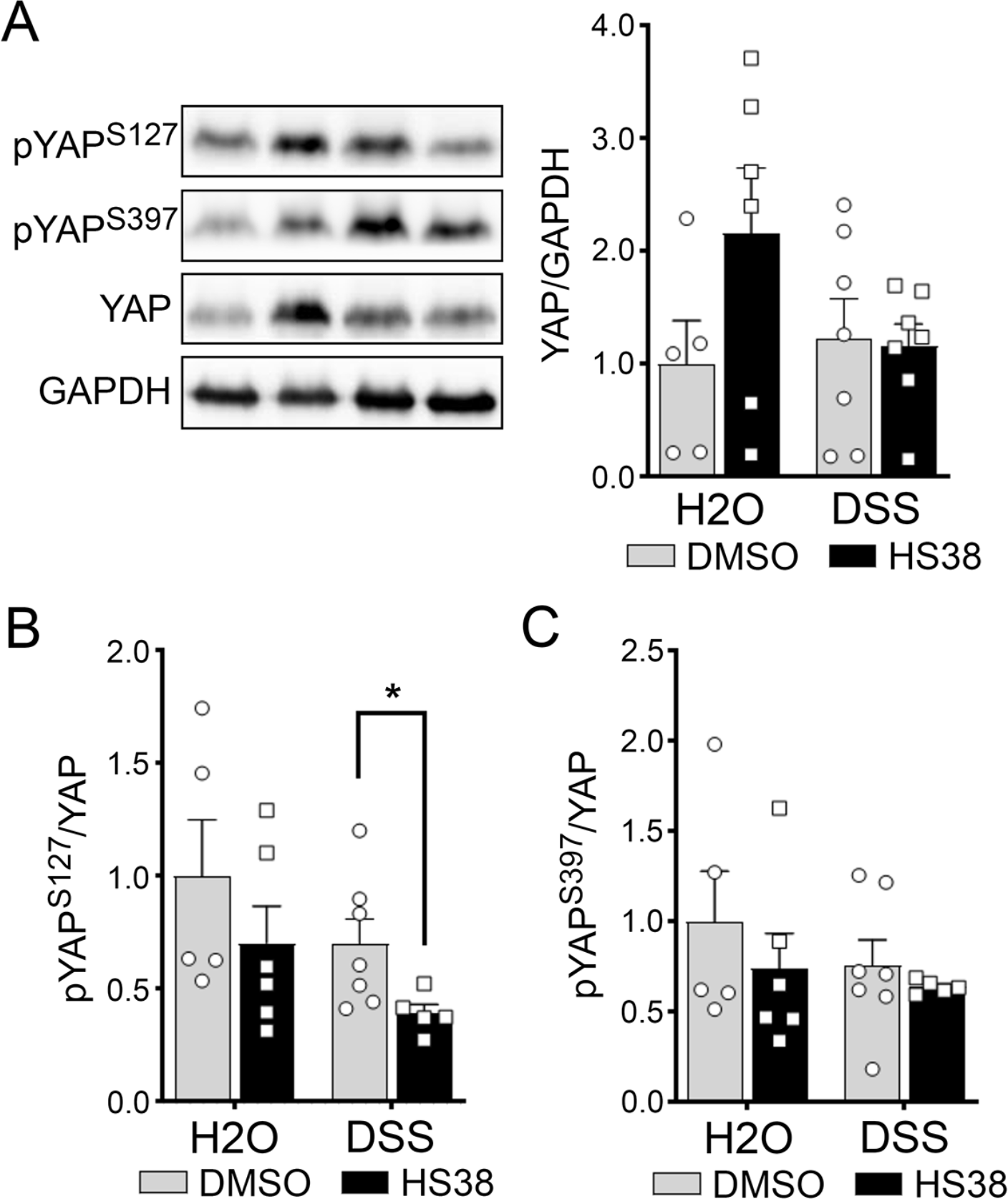
DAPK3 inhibition was associated with a decrease in pYAP-S127 phosphorylation following DSS-induced injury. Tissue lysates obtained from mouse colon midsections were immunoblotted for YAP **(A)**, pS127-YAP **(B)**, and pS397-YAP **(C)**. GAPDH was used as loading control for YAP whereas the pan-YAP signals were used to normalize the pS127-YAP and pS397- YAP levels. Error bars represent ± SEM, n=5-7. Statistical significance between vehicle and HS38 treatments was determined via unpaired Student’s t-test, * p<0.05.

**Figure 9.**
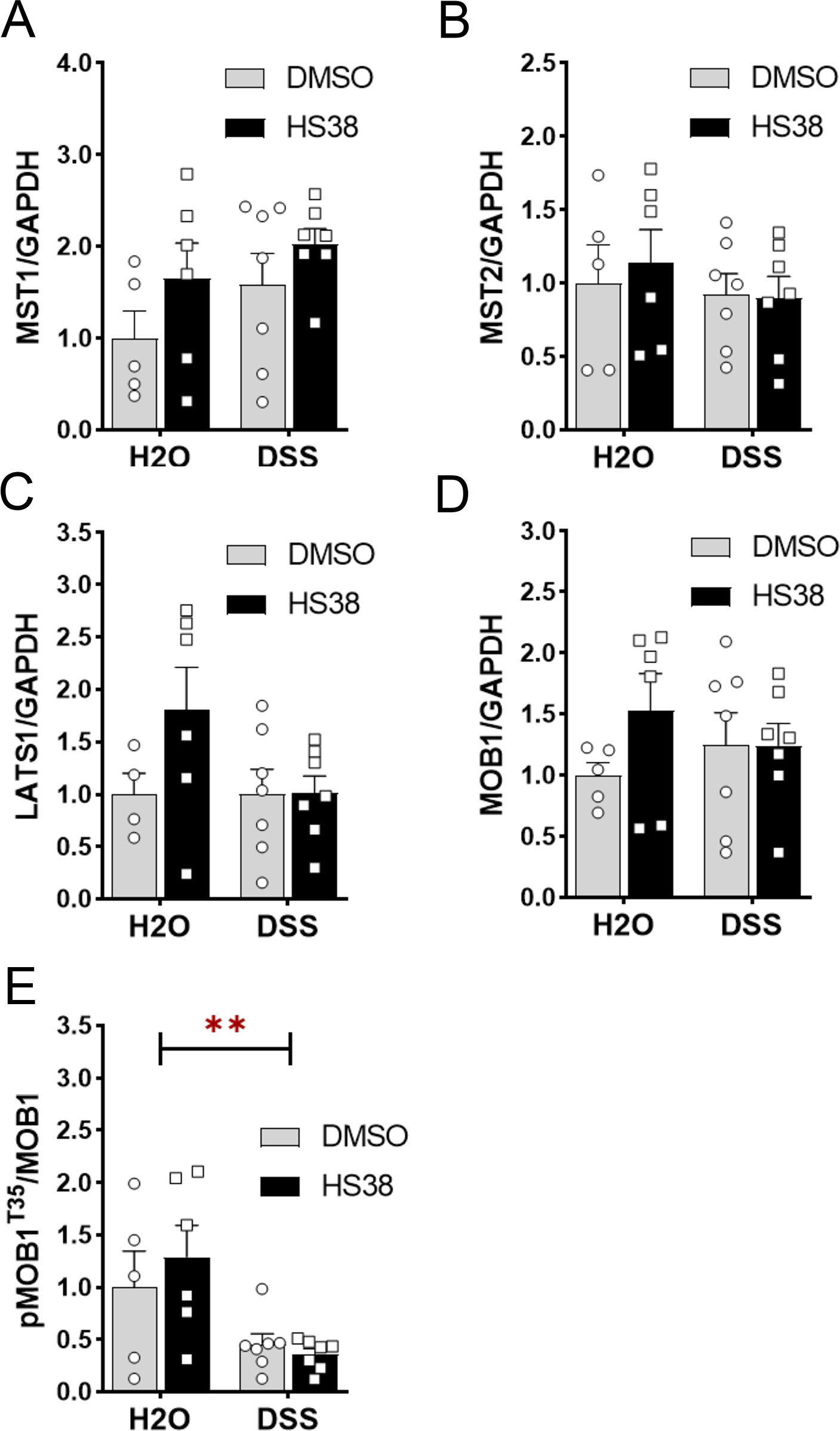
Core Hippo pathway regulators display insignificant changes in protein abundance upon DAPK3 inhibition. Lysates obtained from mice colon midsections were immunoblotted for core regulators of the Hippo pathway, including **(A)** mammalian Sterile 20-like kinases 1 and 2, MST1/2; **(B)**; large tumor suppressor 1 kinase, LATS1, as well as MOB kinase activator 1, MOB1 **(C)** and its regulatory T35 phosphorylation site **(D)**. GAPDH was used as a loading control, but pan-MOB1 was used to normalize the pT35-MOB1 levels. The results represent the mean normalized protein abundance. Error bars represent ± SEM, n=5-7. Statistical significance was determined via unpaired Student’s t-test, ** P<0.01.

The phosphorylation of Ser127 specifically primes the binding of YAP to 14-3-3 and leads to its cytoplasmic retention (Zhao *et al*, 2007). To substantiate the results obtained from pS127- YAP immunoblots, paraffin-embedded distal-colon tissues were probed with antibodies targeting total YAP to evaluate its subcellular localization. In animals receiving both DSS and HS38, colonic IECs showed increased nuclear translocation of YAP when compared to animals receiving DSS and vehicle (**Figure 10A**); this nuclear accumulation of YAP was observed in IECs located at the crypt base and the crypt apex (**Figure 10B**, insets (iii) and (iv) respectively). These data suggest that HS38 treatment, and hence DAPK3 inhibition, bypassed the canonical Hippo pathway to activate YAP.

**Figure 10.**
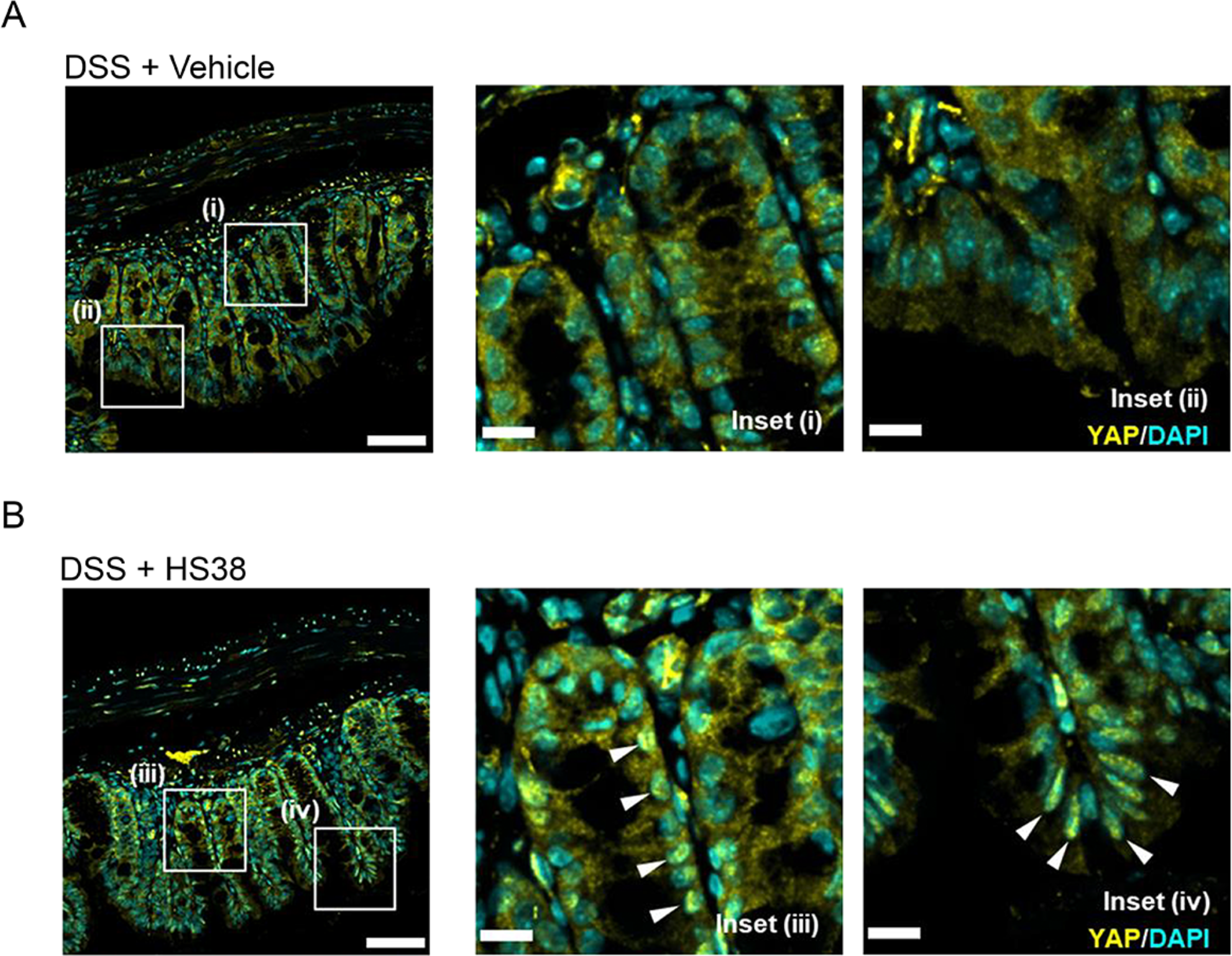
DAPK3 inhibition stimulated YAP nuclear accumulation in colonic IECs following DSS- induced injury. Distal-colon tissue sections obtained from mice treated with DSS + DMSO **(A)** or DSS + HS38 **(B)** were immunostained for YAP (yellow) and nuclei (DAPI, blue). White arrowheads indicate nuclear accumulation of YAP. Scale bars = 100 μm / 20 μm (inset).

## 4. DISCUSSION

Our findings advance DAPK3 as a novel regulator of intestinal epithelial repair and establish a unique linkage between the kinase function of DAPK3 and the Hippo effector YAP within the context of UC. *In vitro* cell studies revealed increased IEC proliferation that underwrote the enhanced wound closure observed in monolayers treated with small molecule inhibitors of DAPK3. Additionally, DAPK3 inhibition promoted colocalization of YAP with F-actin, which is indicative of YAP activation. In mice recovering from DSS-induced UC, the inhibition of DAPK3 also prompted YAP activation. However, DAPK3 inhibition repressed injury-induced IEC hyperproliferation, which elicited worsening of disease severity in the animal model. While this discrepancy may be the consequence of interspecies differences on the regulation of DAPK3 subcellular localization (Weitzel *et al*, 2011), similar effects of DAPK3 inhibition on YAP activation, observed in both Caco-2 IECs and mice, suggests that the conflict is more likely caused by the molecular impact of DAPK3 in non-epithelial cell types, such as macrophages.

The essential role of Hippo signaling on repair from acute intestinal injury has been demonstrated in multiple studies. As the downstream effector of the Hippo pathway, the transcriptional regulator YAP has been implicated as an important mediator of mechanotransduction and proliferation during colonic epithelial repair. With regards to the DSS model of colitis, Cai and colleagues showed impaired epithelial regeneration of *Yap*-deficient colonic crypts (Cai *et al*, 2010). Moreover, Yui and colleagues found that the colonic epithelium undergoes a YAP-dependent transition to fetal-like state to facilitate repair during recovery (Yui *et al*, 2018). Mucosal healing is also crucial for tempering the progression of UC (Boal Carvalho *et al*, 2016), and YAP activity was suggested to promote mucosal regeneration for UC recovery. In this case, IEC-specific YAP ablation in mice increased weight loss and tissue damage with DSS- associated colitis and was shown to impede regeneration of colonic crypts (Cai *et al*, 2010; Taniguchi *et al*, 2015).

DAPK3 may regulate YAP-mediated cell proliferation and cell cycle progression via various molecular pathways. For one, DAPK3 may exert control on YAP activity through Wnt signaling, as DAPK3 knockdown was shown to reduce β-catenin/TCF4 mediated-transcriptional activities (Togi *et al*., 2011), and Wnt signaling was demonstrated to induce YAP activity during crypt regeneration (Guillermin *et al*, 2021). However, it should be noted that DAPK3 kinase activity was not required for its regulation of Wnt signaling (Togi *et al*, 2011), and Wnt signaling could induce YAP activity via increased transcription of *YAP* rather than through post-translational regulation of YAP protein localization (Guillermin *et al*, 2021). Thus, the impact of DAPK3 on YAP, brought about through HS38 administration, is unlikely to be facilitated via Wnt signaling.

Additionally, DAPK3 may promote YAP activity through mechanical cues, elicited by changes to cell tension that occur with the phosphorylation of myosin regulatory light chain (MLC20). Actomyosin, composed of F-actin and non-muscle myosin II, serves as a central mediator between mechanical cues and Hippo/YAP signaling (Nardone *et al*, 2017). DAPK3 can directly phosphorylate MLC20 (Komatsu & Ikebe, 2004; Moffat *et al*, 2011) and may also influence actomyosin-dependent cellular tension through its inhibition of myosin light chain phosphatase (MLCP) (MacDonald *et al*, 2001). Merlin/NF2, an upstream regulator of the Hippo pathway, was demonstrated to be a substrate of MLCP (Jin *et al*, 2006), and recent evidence showed that the knockdown of *MYPT1* (i.e., PPP1R12A, the myosin phosphatase-targeting subunit of MLCP) in ovarian cancer cell lines was associated with increased Merlin/NF2 phosphorylation and coincident activation of the Hippo pathway (Munoz-Galvan *et al*, 2020). As DAPK3 inhibition with HS38 ought to enhance MLCP function, thus strengthening the activation of Merlin/NF2 to reduce YAP activity, the (de)regulation of MLCP function via DAPK3 inhibition does not explain the outcomes observed in the cell studies. However, the substrate preferences of DAPK3 within the locational context of the IEC intracellular environment remain to be described.

In terms of IEC proliferation, no changes were observed between HS38-treatment and vehicle control in mice at baseline. However, in >50% of uninjured mice (i.e., mice receiving H2O instead of DSS), YAP protein abundance increased in response to administration of HS38. As well, a trend was found towards a reduction in phosphorylation of YAP on S127 in these animals. Given that YAP is largely dispensable in homeostasis, it was not surprising to find a lack of prominent defects in these control animals despite the alteration of YAP protein abundance and YAP activation by HS38. Unexpectedly, however, the hyperproliferative phenotype of Caco-2 cells treated with HS38 was not observed in the DSS-model of animal colitis. Instead, animals treated with HS38 displayed a significant decrease in IEC proliferation. This anti-proliferative effect of HS38 on IECs likely played a part in the worsening severity of DSS-induced colitis. Paradoxically, in mice recovering from DSS-induced injuries, pS127-YAP levels decreased significantly in the HS38 treatment group. Further, DAPK3 inhibition stimulated YAP nuclear accumulation in colonic IECs. These data suggest that DAPK3 inhibition prompted YAP activation, which ought to have enhanced IEC regeneration. One plausible explanation for such paradoxical phenotype is that the IEC proliferation in mice recovering from DSS-induced injury was depressed by the functioning of YAP in other cell types.

A recent study suggests that YAP can drive macrophages toward M1 polarization while restricting M2 polarization during UC development (Zhou *et al*, 2019). While IEC proliferation was not specifically examined, myeloid-specific ablation of YAP could attenuate the severity of DSS- induced colitis; YAP impeded M2 polarization, promoted M1 polarization, and increased production of IL-6 (Zhou *et al*, 2019). Given that IL-6 signaling can trigger YAP activation via stimulation of gp130-associated Src family kinase Yes (Taniguchi *et al*, 2015), a positive feedback loop between YAP and IL-6 production by macrophages likely exists. Potentially, DAPK3 inhibition brought about YAP activation in macrophages, which could impair M2 polarization to depress IEC proliferation post-DSS-induced injury. The regulation of macrophage programming by DAPK3 was also demonstrated (Mukhopadhyay *et al*, 2008); DAPK3 restrained IFN-γ induced gene expression in macrophages through its phosphorylation of the ribosomal protein L13a. The phosphorylation of L13a activated the IFN-γ-activated inhibitor of translation complex to repress translation of inflammatory genes (Mukhopadhyay *et al*, 2008). Taken together, DAPK3 may contribute an important role in the macrophage resolution of inflammation program and take part in a yet undefined counteracting mechanism that resets the positive feedback loop between YAP and IL- 6 in macrophages.

Ultimately, a genetic approach will likely be necessary to dissect the independent roles that DAPK1 and DAPK3 play in intestinal epithelial repair and Hippo signaling. However, the current absence of genetic resources precludes the confirmation of results obtained with pharmacologic inhibition of DAPK3 using mouse models. Global *Dapk3* knockout is embryonically lethal for mice (Kocher *et al*, 2015), with similar lethality observed for endothelial-cell specific *Dapk3* knockout (Zhang *et al*, 2019). While inducible-*Dapk3* whole-body knockout and smooth-muscle specific *Dapk3* knockout mice were successfully generated (Zhang *et al*, 2019), *Dapk3* knockout cell lines created with CRISPR-Cas9 gene editing could not be maintained, suggesting an essential role for DAPK3 in cell growth and/or survival (Takahashi *et al*, 2021).

An advantage of small molecule inhibitors is that they can be titrated to reveal a spectrum of phenotypes. Moreover, the inhibition of enzymatic activity does not necessarily prevent protein-protein interactions. Therefore, the use of small molecule inhibitors presents an opportunity to differentiate between the enzymatic and structural roles of proteins. Several uncommon features in the ATP-binding pocket of DAPK3 underwrite the high selectivity offered by HS38; nevertheless, HS38 still displays high potency towards DAPK1 and the structurally related PIM3 kinase (Carlson *et al*, 2013). So, it is necessary to consider the off target impacts of DAPK1 and PIM3 on IEC wound healing when utilizing HS38. The PIM3-mediated contribution of HS38 to wound healing was addressed in this study (**Figure 2**). Parallel *in vitro* experiments completed by treatment of Caco-2 cells with HS38, HS56 or HS94 facilitated the assignment of observed effects as being DAPK3-centric or PIM3-centric. Unfortunately, with regards to DAPK1, all currently available DAPK3 inhibitors demonstrate comparable potency towards both DAPK1 and DAPK3 (Al-Ghabkari *et al*, 2016; Carlson *et al*, 2013; Okamoto *et al*, 2010). Thus, the crosstalk between DAPK3 and DAPK1 and the potential for overlapping biological functions will need to be assessed in future studies. This may be possible once additional structure-activity relationship (SAR) studies inform the construction of pharmacophores that can differentiate between DAPK3 and DAPK1.

## ACKNOWLEDGEMENTS

This work was supported by a research grant from the Canadian Institutes of Health Research (MOP#97931 to J.A.M.). H.-M.C. was recipient of CIHR Fredrick Banting and Charles Best Canada and Alberta Graduate Excellence Scholarships.

## 5. AUTHOR CONTRIBUTIONS

H.-M.C. completed the data analysis, prepared figures, and wrote the manuscript. D.A.C synthesized the DAPK3 inhibitor compounds HS-38, HS-94 and HS-56. T.A.J.H. coordinated inhibitor compound production and made intellectual contributions to the project. J.A.M. conceived and coordinated the study, assisted with experimental design, wrote the manuscript, provided trainee supervision, and made intellectual contributions to the project. All authors reviewed the results and approved the final version of the manuscript.

